# Two waves of evolution in the rodent pregnancy-specific glycoprotein (PSG) gene family lead to structurally diverse PSGs

**DOI:** 10.1101/2023.02.21.529360

**Authors:** Robert Kammerer, Wolfgang Zimmermann

## Abstract

The evolution of pregnancy-specific glycoproteins (PSGs) within the CEA gene family of primates correlates with the evolution of hemochorial placentation about 45 Myr ago. Thus, we hypothesized that hemochorial placentation with intimate contact between fetal cells and maternal immune cells favors the evolution and expansion of PSGs. With only a few exceptions, all rodents have hemochorial placentas thus the question arises whether PSGs evolved in most rodent genera.

Analyzing genomic data of 94 rodent species we could identify PSGs only in three families of the suborder Myomorpha (characteristic species in brackets) namely in the Muridae (mouse), Cricetidae (hamster) and Nesomyidae (giant pouched rat) families. No PSGs were detected in the suborders Anomaluromorpha (springhare), Castorimorpha (beaver), Hystricognatha (guinea pig) and Sciuromorpha (squirrel). Thus, PSGs evolved only recently in Myomorpha shortly upon their most recent common ancestor (MRCA) has coopted the retroviral genes syncytin-A and syncytin-B which enabled the evolution of the three-layered trophoblast. This may suggest that the evolution of *Psgs* in rodents may have been favored by the challenge of the newly invented architecture of the maternal-fetal interface. In addition, a second hallmark of rodent PSG evolution seems to be the translocation of genes from the CEA gene family locus into a unique genomic region. Rodents without PSGs do not have any CEA-related genes in this locus. In contrast, rodent species in which PSGs evolved have lost ITAM-encoding CEACAM genes indicating that such a gene was translocated and thereby destroyed to form the new rodent PSG locus. This locus contains at least one *Psg* and *Ceacam9* indicating that one of them was the founder gene of rodent *Psgs*. These genes are composed of various numbers of IgV-like domains (N domains) and one carboxy-terminal IgC-like domain of the A2-type. In a second wave of gene amplification in the PSG locus a gene encoding a protein composed of two N domain gave rise to four genes in mice (*Ceacam11*-*14*). In light of the divergent structure of PSGs in various mammalian species, we hypothesized that the *Ceacam11-14* encode also functional PSGs and indeed we found that they are preferentially expressed by spongiotrophoblast cells, like *Psg* genes.

## Background

Pregnancy-specific glycoproteins (PSGs) were first described in humans as proteins in the serum of pregnant women [1]. Subsequently, the genes which encode human PSGs were identified and found to be members of the carcinoembryonic antigen (CEA) gene family which by itself is a member of the immunoglobulin gene superfamily [2, 3]. Once the CEA gene families were investigated in mice and rats a subgroup of the gene products was identified as secreted glycoproteins that were predominantly expressed by trophoblast cells and were named as PSGs in rodents [4-6]. Surprisingly, the structure of rodent PSGs differs significantly from that of human PSGs [6]. While human PSGs are composed of one N terminal immunoglobulin variable (IgV)-like domain (also called, N domain) and two to three immunoglobulin constant (IgC)-like domains (two A and one B domains) murine PSGs contain three to eight N domains followed by a single IgC domain of the A2-type found among others in CEACAM1 [7]. This led to the assumption that primate and rodent PSGs evolved independently in both orders. More recently, we found that in some microbat species, putative PSGs exist, composed of a single N domain followed by a single A domain [8]. Furthermore, in horses PSGs consisting of a single N domain were identified [9]. Despite the vast structural differences, common functions were described for PSGs of different species such as inhibition of platelet aggregation, activation of latent TGFβ and other immune-modulating functions [9-13] suggesting that PSGs developed independently in different mammalian lineages by convergent evolution [14]. This raises the question about the driving force of PSG evolution within the CEA gene family. Based on the fact that humans, mice, and rats as well as the above-indicated bat species have a hemochorial placenta, where fetal trophoblast cells have direct contact with maternal immune cells we and others hypothesized that PSGs evolved to regulate maternal immunity against fetal antigens [15, 16]. Indeed, it was found that equine PSGs were expressed by highly invasive trophoblast cells the so-called girdle cells which later form endometrial cups, a unique structure in equine placenta [9]. It is well documented that these cells are recognized by the maternal immune system which is also expected for trophoblast cells in mammals with hemochorial placentation [17]. Furthermore, in primates PSGs were found only in species with hemochorial placentas but not in primates that have an epitheliochorial placenta further pointing to an association of PSG evolution and intimate interaction of fetal trophoblast cells and the maternal immune cells [16]. Rodents, with only very few exceptions, have a hemochorial placenta, so we wondered when the PSGs evolved in rodents [18]. Rodents first appear in the fossil record at the end of the Paleocene and earliest Eocene, about 54 million years ago (Mya) [19]. Nowadays, the order Rodentia comprises about 40 % of all mammalian species [20] and is divided into five suborders, the Anomaluromorpha (e.g. springhares), Castorimorpha (e.g. beavers and kangaroo rats), Myomorpha (e.g. mice and hamsters), Hystricomorpha (e.g. guinea pigs and chinchillas) and the Sciuromorpha (e.g. squirrels and mountain beavers) [21]. Mice and Rats belong to the Myomorpha suborder which appeared ~26 mya. The *Mus-Rattus* split is estimated to have occurred 8.8 to 10.3 mya ago [22]. Since PSGs in mice and rats are thought to have a common ancestor this indicates that PSGs in rodents evolved at least about 10 mya ago. But what happened during the remaining 40 million years of rodent existence? To answer this question, we investigated the CEA gene families in 94 rodent species containing members of the rodent suborders Myomorpha, Hystricomorpha, Sciuromorpha, and Castorimorpha. We found only supporting evidence for the evolution of PSGs in Muroidea, a subgroup of the Myomorpha, not in other rodents. The key event for the amplification of PSGs was most likely the translocation of CEA gene family member(s) or parts of them from the CEA gene family locus into the *Npas1*/*Pglyrp1* locus. In this locus three, structurally different members of the CEA gene family could be found which all encode secreted glycoproteins. PSGs consists of multiple N domains and a single A domain, Ceacam11-14 consists of two N domains, and Cecam9 and Ceacam15 are composed of one N domain and one A domain. According to their expression pattern in mice, all of them have to be considered to be functional PSGs. Thus, domain arrangements of PSGs do not only differ fundamentally between species but also within a single species.

## Results

### Recent evolution of pregnancy-specific glycoproteins in rodents

*Psgs* are well described for mice and rats but so far not for other rodents. In mice and rats, *Psgs* are located in the genome locus flanked by marker genes *Npas1* and *Pglyrp1* [23]. This locus will be further referred to as the “rodent *Psg* locus” in this publication. In contrast, no CEA gene family members are present in this region in primate genomes [16]. In mice, in addition to the 17 *Psgs* (*Psg16*-*Psg32*) *Ceacam9* and *Ceacam11-15* are located at this locus [7, 24]. To get first insights into the evolution of *Psgs* in rodents, other than mice and rats, we used the sequences of the above-mentioned mouse genes to identify *Psgs* in the genome of 94 rodent species using the Basic Local Alignment Search Tool (BLAST) and the NCBI and Ensemble databases (Fig. 1, Supplementary Table 1). *Psgs*, composed of three or more N domains and one IgC-like domain as described for mice and rats, were identified only in the suborder Myomorpha but not in the suborders Castorimorpha, Hystricomorpha and Sciuromorpha (Fig. 1).

**Figure 1.**
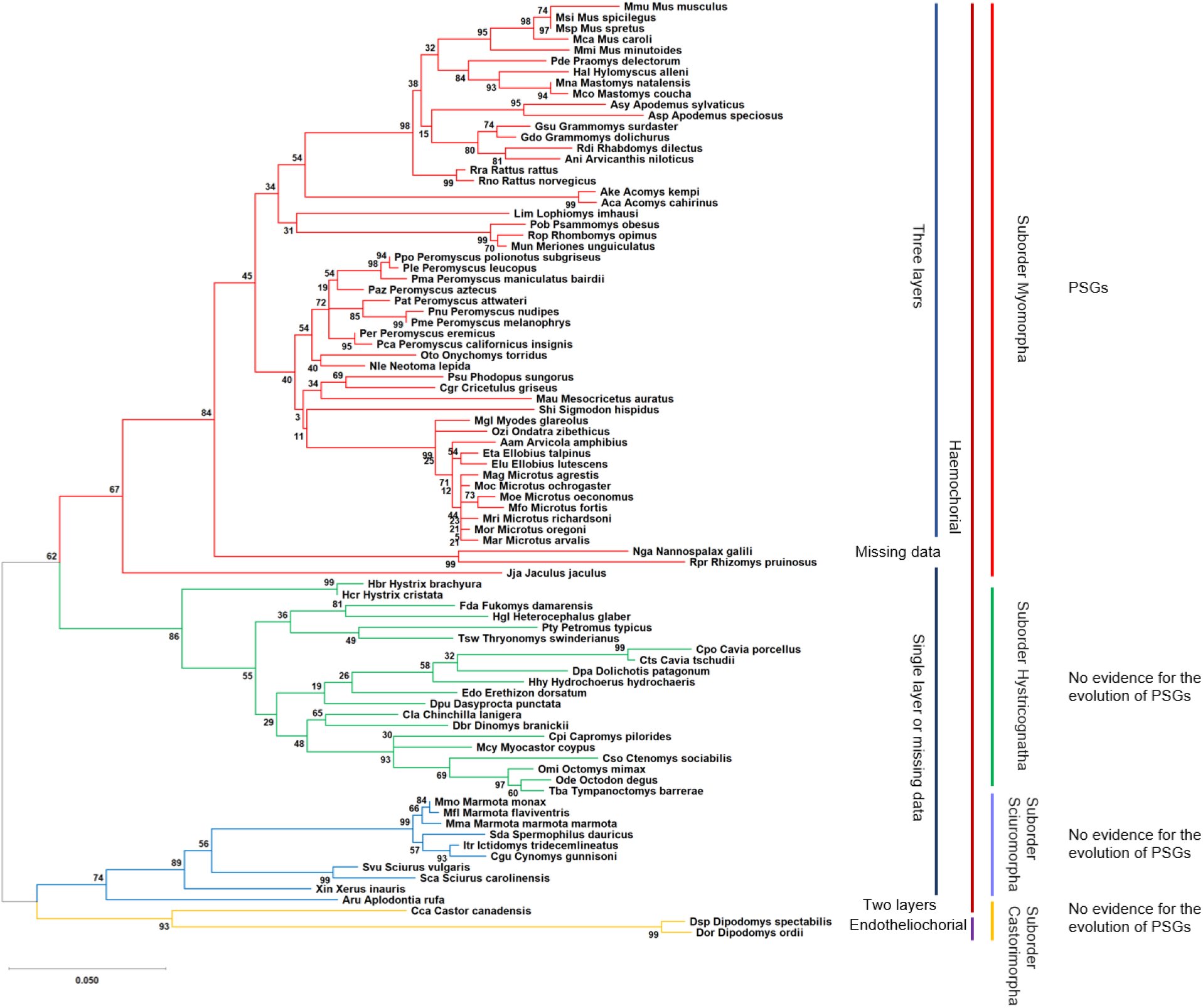
Phylogeny of the analyzed rodent species. The phylogenetic tree was constructed using the N domain exon nucleotide sequences from *Ceacam19* genes. The color of the branches indicates the suborder to which the species belong. The type of placenta is indicated on the right. The presence of *Psg* genes is indicated.

*Psgs* were found in all analyzed species of the suborder Myomorpha except in the genome of the lesser Egyptian jerboa (Jaculus jaculus; Dipoditae) and the two members of the Spalacidae family the Upper Galilee mountains blind mole rat (*Nannospalax galili*) and the hoary bamboo rat (*Rhizomys pruinosus*) (Fig. 2). Thus, the presence of *Psgs* is limited to three rodent families (Cricetidae, Muridae and Nesomyidae) of the Muroidea clade. Interestingly, the number of *Psgs* varied widely from three genes in the genome of the African giant pouch rat (*Cricetomys* gambianus) a member of the Nesomyidae family and of the Mongolian gerbil (*Meriones unguiculatus*), the great gerbil (*Rhombomys opimus*) and the fat sand rat (*Psammomys obesus*) all three are members of the Gerbillinae subfamily, to 25 genes (including 2 pseudogenes) in the North American deer mouse (*Peromyscus maniculatus*; Neotominae subfamily) (Fig. 2). Interestingly, in all rodent species where we identified *Psgs* we also identified *Ceacam9* orthologs, although in the three Gerbillinae species (*Meriones unguiculatus, Rhombomys opimus, Psammomys obesus*) *Ceacam9* seem to be a pseudogene due to a common two nucleotide deletion in the N domain exon (Fig. 2). *Ceacam15* orthologs were found in all species which have *Psgs* and *Ceacam9* except in species of the Arvicolinae subfamily (Fig. 1, Fig. 2). However, a possible remnant of *Ceacam15* was found in the two Ellobius species as well as in the genome of *M. glareolus* and *O. zibethicus*, indicating that *Ceacam15* was lost in the Arvicolinae subfamily. In species that do not have *Psg* genes, neither *Ceacam9* orthologs nor *Ceacam15* orthologs were found (Fig. 2). Genes related to murine *Ceacam11-14* are found in a subgroup of the species with *Psgs* and are described in more detail below.

**Figure 2.**
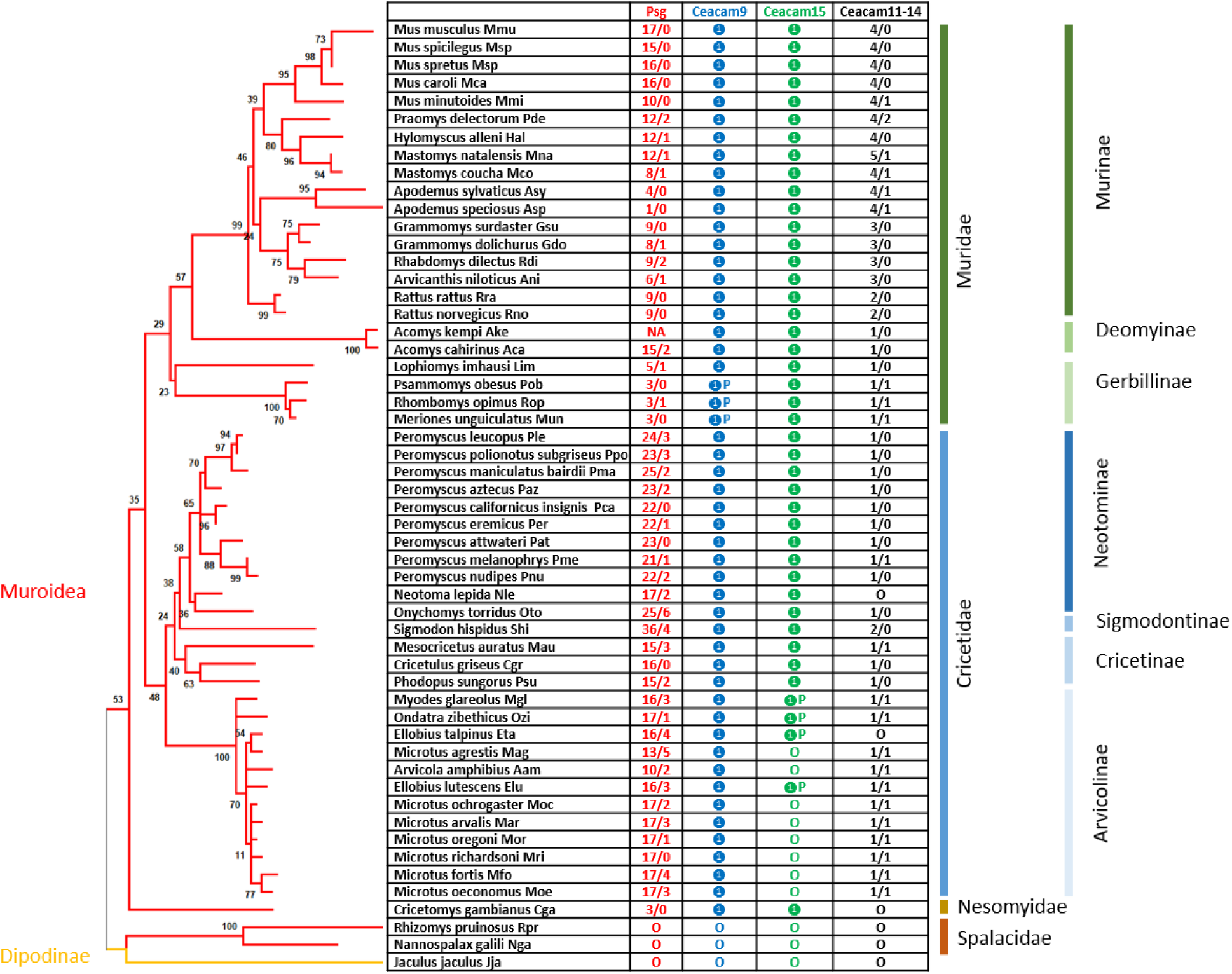
PSGs evolved in the clade Muroidea. The phylogenetic tree was constructed with the nucleotide sequences of the *Ceacam19* N domain exons. Genes found in the genome locus flanked by marker genes *Npas1* and *Pglyrp1* were analysesd in 54 species of the suborder Myomorpha. Open circles indicate that the gene was not found in the genome; filled circles specify that the gene was identified in the genome as a single copy; P next to the filled circle indicates that the gene is a pseudogene according to our definition (Material and Methods). NA = not analysed; The numbers indicate the total number of genes identified/the number of genes expected to represent pseudogenes.

### Coincidence of *Psg* appearance at the “rodent *Psg* locus” and loss of ITAM-encoding *Ceacams* in rodent CEA gene families

In order to get further information about the possible origin of *Ceacam*-related genes at the rodent *Psg* locus we analysed the chromosomal arrangement of *Ceacam*-related genes. The analyses were restricted to species for which available scaffolds were long enough to cover the entire *Ceacam*/*Psg* locus. Selected species are depicted in Fig. 3. Remakably, species which lack *Psgs* do also not harbor any other members of the *Cea* gene family in the “rodent *Psg* locus” (Fig. 3). This was verified for species belonging to the Suborders Hystricomorpha and Sciuromorpha as well as to the members of the Spalacidae family (Fig. 3). This may indicate that a single translocation of one or more Cea gene family members gave rise to the evolution of all Cea gene family members in the “rodent *Psg* locus”. However, in the *Ceacam* locus diverse differencies and copy number variations could be observed. Interestingly, we observed that rodent species which do not have Cea gene family members in the “rodent Psg locus” have Cea gene family members encoding Ceacams which have activating singnaling motifs in the cytoplasmic tails (Supplementary file 1). Of note such Ceacams were not found in rodent species in which *Psgs* evolved including mice and rats. In some species e.g. the alpine marmot (*Marmota marmota*) such genes even have been multiplied (Fig. 3, Supplementary file 1). This may tempt to speculate that an activating *Ceacam* was destroyed and subsequently lost due to the translocation of a *Ceacam* gene to form the “rodent Psg locus” in the MRA of *Psg* harboring rodents.

**Figure 3.**
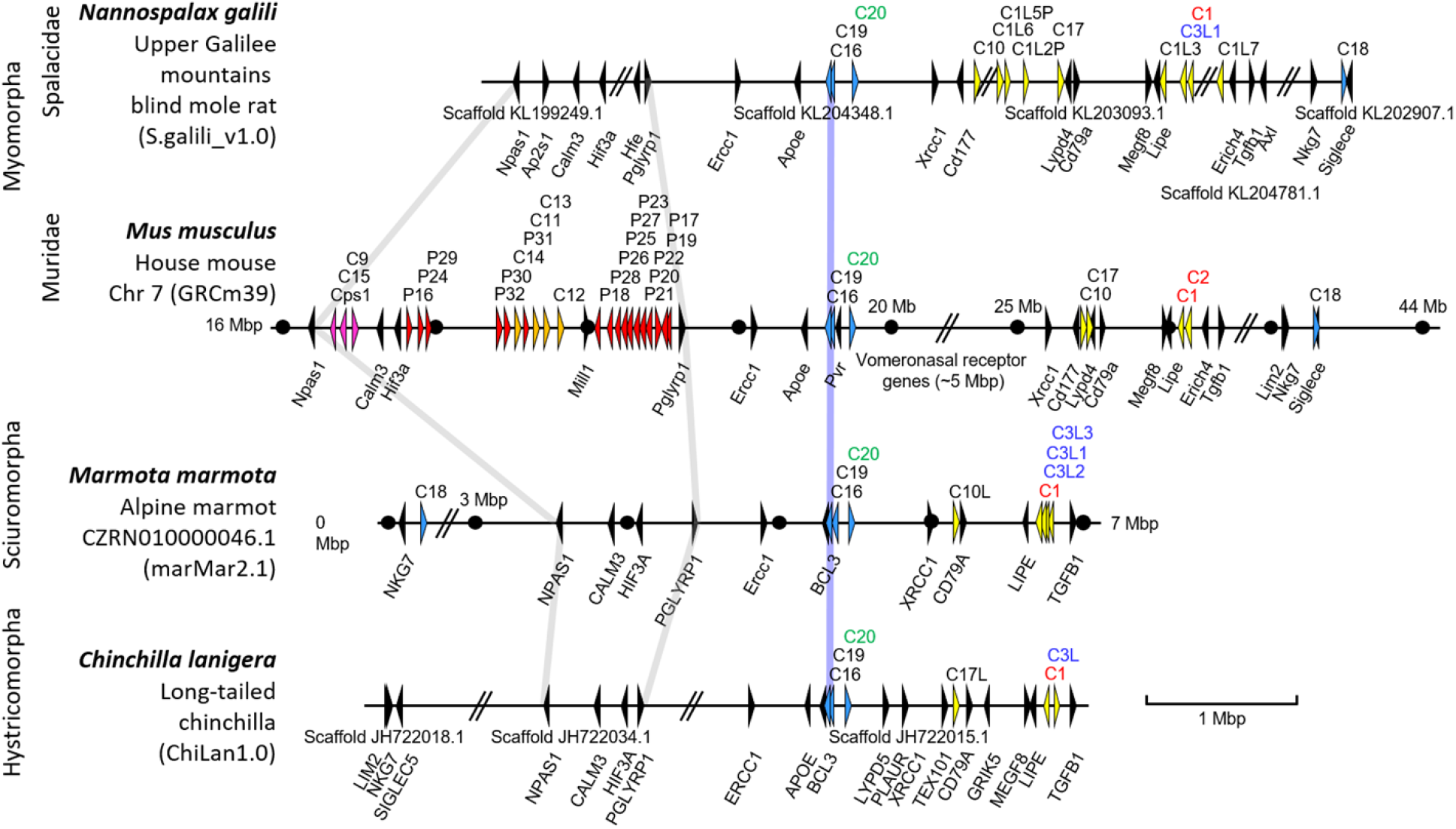
Evolution of *Psgs* in rodents is restricted to the *Npas1*/*Pglyrp1* locus. The chromosomal arrangements of *Ceacam*-related genes of selected species of the Myomorpha (*Nannospalax galili, Mus musculus*), Sciuromorpha (*Marmota marmota*), and Hystricomorpha (*Chinchilla lanigera*) suborders are shown. Arrowheads indicate genes with their transcriptional orientation. The *Psg*-related genes are shown in red (*Psg*), purple (*Ceacam9, Ceacam15*, and Ceacam pseudogene 1, *Cps1*) or orange (*Ceacam11-14*), *Ceacam1*-related Ceacam genes in yellow, conserved *Ceacam* genes in blue and selected flanking genes in black. The *Ceacam* gene loci were aligned along the position of *Ceacam16* (blue line). Gray lines were used to delineate the *Npas1*/*Pglyrp1* loci. Abbreviated names of *Ceacam1*-like genes with ITIM/ITSM-encoding exons are shown in red and with ITAM and ITAM-like motif-encoding exons in green and blue, respectively. Nucleotide numbering of the chromosomes starts at the telomere. Selected positions 1 Mbp apart are indicated by dots. Databases and their versions used are listed below the species name. The borders of scaffolds are indicated by double slashes, their names below the chromosome. The exact distances between the scaffolds still have to be determined by complete whole genome sequencing. Of note: The rodent genomes (except the murine genome) are not completely sequenced yet. Therefore, not all *Ceacam* genes identified in whole genome shotgun (WGS) databases have been found in the published assembled genomes. C, *Ceacam*; Cps, *Ceacam* pseudogene; C1L1(P), Ceacam1-like (pseudo)gene, the same abbreviation schema applies to similar abbreviations; Mbp, million base pairs; P, pregnancy-specific glycoprotein (*Psg*) genes.

### A second wave of gene amplification led to the generation of murine *Ceacam11-14* genes

To further delineate the evolution of the *Ceacam*-related genes at the “rodent *Psg* locus” we performed phylogenetic analyses of N domain exons of members of different muroid families i.e. house mouse and Chinese hamster, using their nucleotide sequences. An orthologous relationship was found for *Ceacam9, Ceacam15, Ceacam16, Ceacam17* and *Ceacam19* (Fig. 4). Furthermore, mouse *Ceacam1, Ceacam2* and *Ceacam10N1* are closely related with *Ceacam1* and *Ceacam2* in the Chinese hamster (*Cricetulus griseus*) but did not exhibit pairwise orthology. For *Psgs* the N1 domain exon sequences build a cluster but no orthologous relationship between individual *Psgs* of the two species could be identified. The N2 and N3-6 N domains did not segregate completely into individual clusters indicating that recent exon duplication and shuffling has taken place during expansion of *Psgs*. Remarkably, in the consensus tree the *Ceacam9* N domain exon is closely related to the N1 domain exons of *Ceacam11-14* in mice and to a *Ceacam11*-like gene in the hamster. In addition, the N2 sequence of murine *Ceacam11-14* cluster together with the N2 domain of the *Ceacam11*-like gene in the hamster. However, hamster C11-like exons N1 and N2 do not exhibit clear orthology to any of the Ceacam11-14 genes. Together, this indicates that murine *Ceacam11-14* genes and the hamster *Ceacam11*-like gene have a common ancestor (Fig. 4).

**Figure 4.**
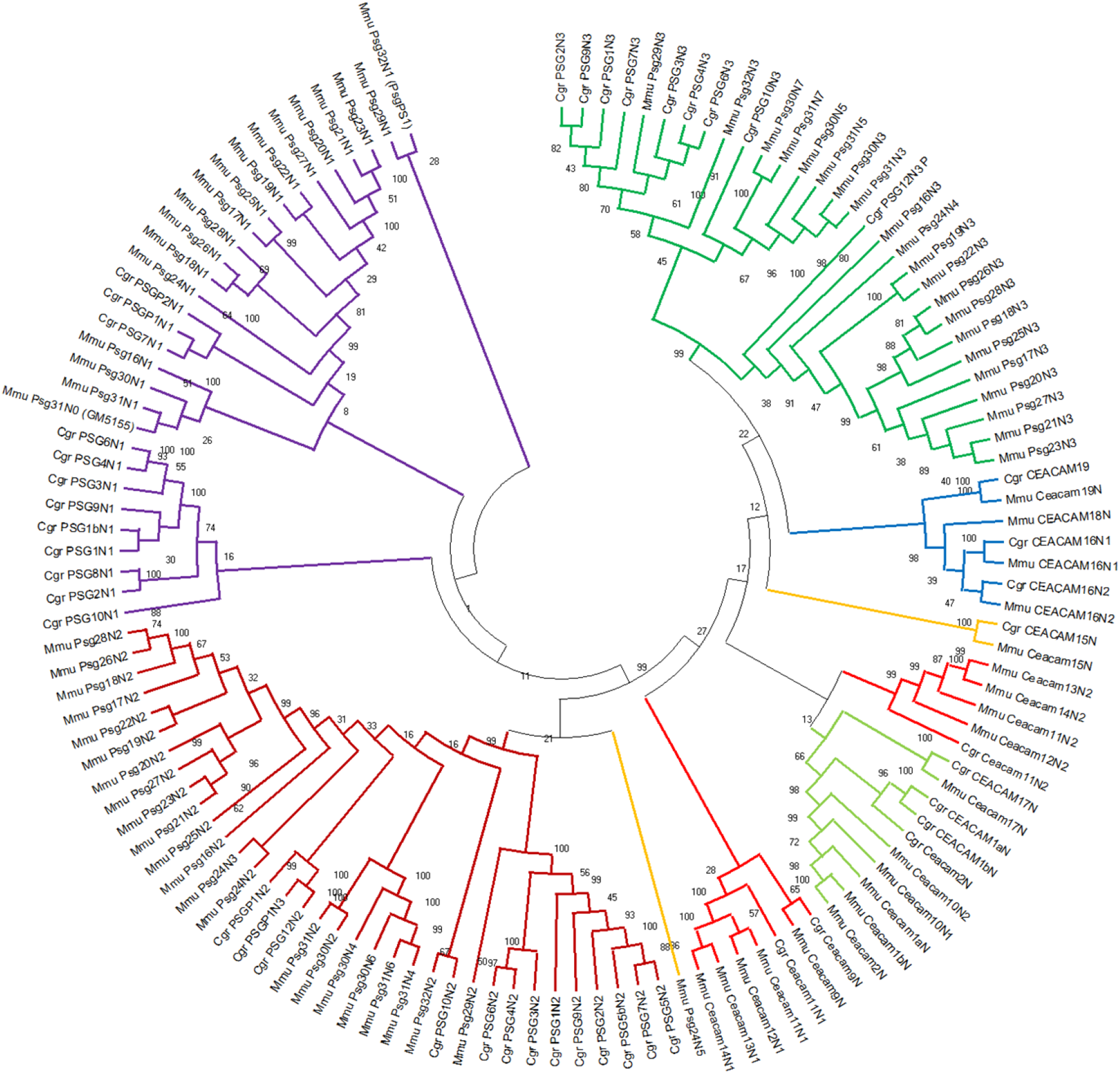
Phylogenetic tree of *Ceacam*/*Psg*-related N domain nucleotide sequences of *Mus musculus* (house mouse) and *Cricetulus griseus* (Chinese hamster). The phylogenetic tree was constructed using the maximum likelihood (ML) method with bootstrap testing (500 replicates). The bootstrap consensus tree inferred from 500 replicates is taken to represent the evolutionary history of the exons analyzed. The percentage of replicate trees in which the associated exons clustered together in the bootstrap test is shown next to the branches. Initial tree(s) for the heuristic search were obtained automatically by applying Neighbor-Join and BioNJ algorithms to a matrix of pairwise distances estimated using the Maximum Composite Likelihood (MCL) approach, and then selecting the topology with superior log likelihood value. Multi-alignment of N domain exon sequences was performed using Muscle implemented in MEGAX. For murine *Psg31* and *Psg32* the name which is currently annotated in the NCBI and Ensemble databases are indicated in brackets. Three letter code abbreviation for species: Mmu, *Mus musculus*; Cgr, *Cricetulus griseus*.

Therefore, we used the nucleotide sequences of the *Ceacam11*-like gene in the hamster to search for closely related N domain exons in other rodent species. With a few exceptions, we identified one to four *Ceacam11*-like genes (composed of two N domain exons) in all species that also have *Psg* and *Ceacam9* genes. A single *Ceacam11*-like gene was found in species of the Cricetidae, Neotominae, and Deomyinae rodent subfamilies. In Murinae an amplification of the *Ceacam11*-like gene had occurred, leading to two genes in rats, three genes in Grammomys, Arvicanthis, and Mastomys, and four genes in the Mus genus (Fig. 2, Fig. 5). In Arvicolinae, only *Ceacam11*-like gene remnants (N2 exons) could be identified. This indicates that this gene was lost in Arvicolinae. Like in the *Psg* genes, orthologous relationship can only be observed in closely related rodent species.

**Figure 5:**
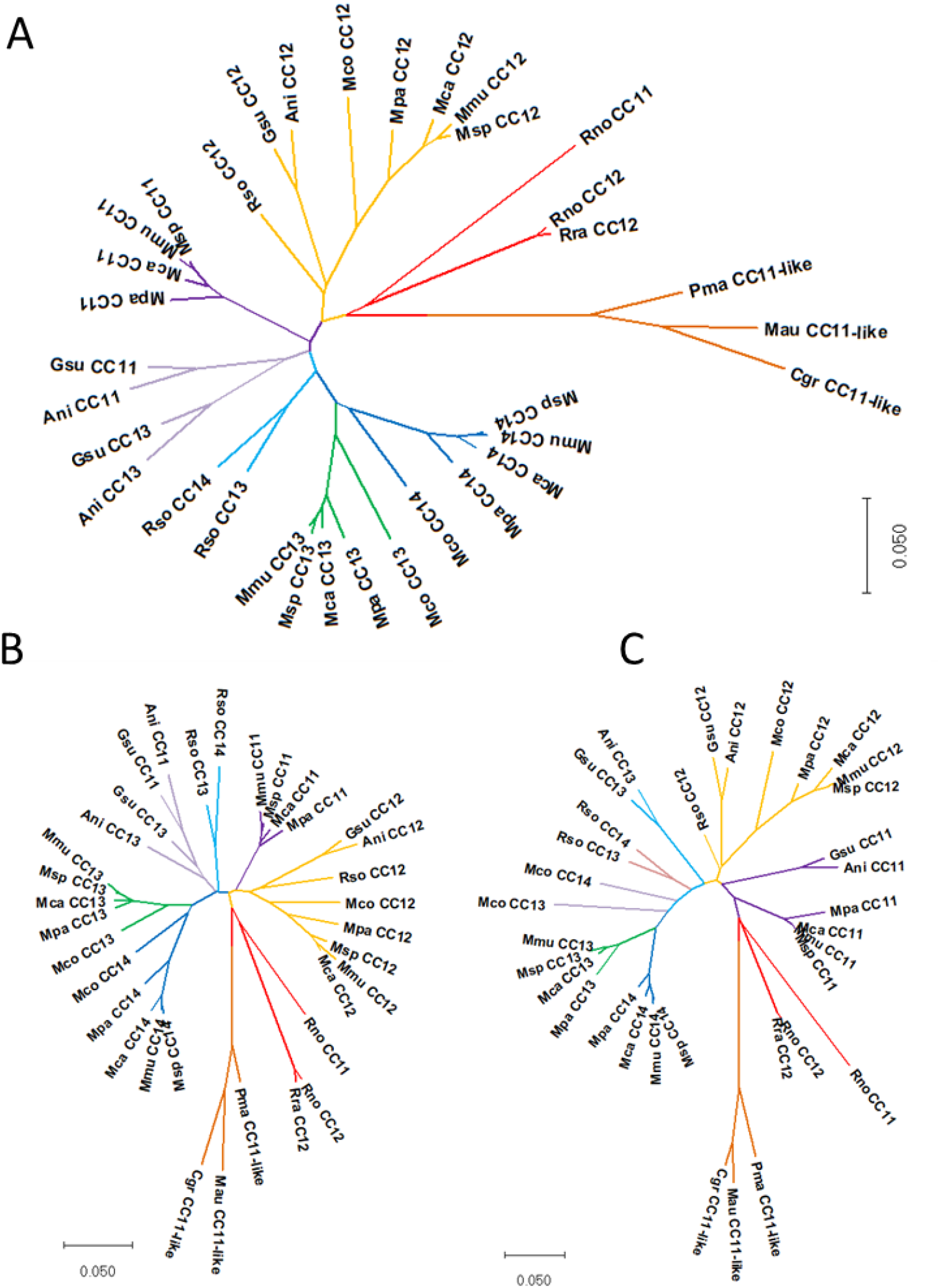
Phylogeny of the *Ceacam11-14* genes in rodents. Nucleotide sequences complete coding sequence (A) the N1 exon (B) and the N2 exon (C) of the *Ceacam11-14* genes were used for phylogenetic analysis. The species abbreviations are explained in Fig. 1 and Supplementary Table 1. CC, Ceacam; P, indicates a missing open reading frame.

### Structure of rodent CEACAM11-14

*Ceacam11-14* genes are in general composed of four exons with encode the leader sequence, the N1 domain and the N2 domain and a 3’ exon harboring the stop codon. Murine *Ceacam14* has a mutation in the splice donor site of exon 3 leading to the usage of a stop codon immediately after the splice donor site. Interestingly, we could not identify exon 4 from rodents with only 1 *Ceacam11*-like gene, however, the splice donor site of exon 3 is intact. Structurally, *Ceacam12* is the most remarkable since the domain encoded by exon 4 is predicted to be part of the ligand binding face of domain N2 which is formed by one of the two β-sheets present in IgV-like domains. This structure is well conserved between different species, indicating that there may exist a common ligand for CEACAM12 (Fig. 6).

**Figure 6:**
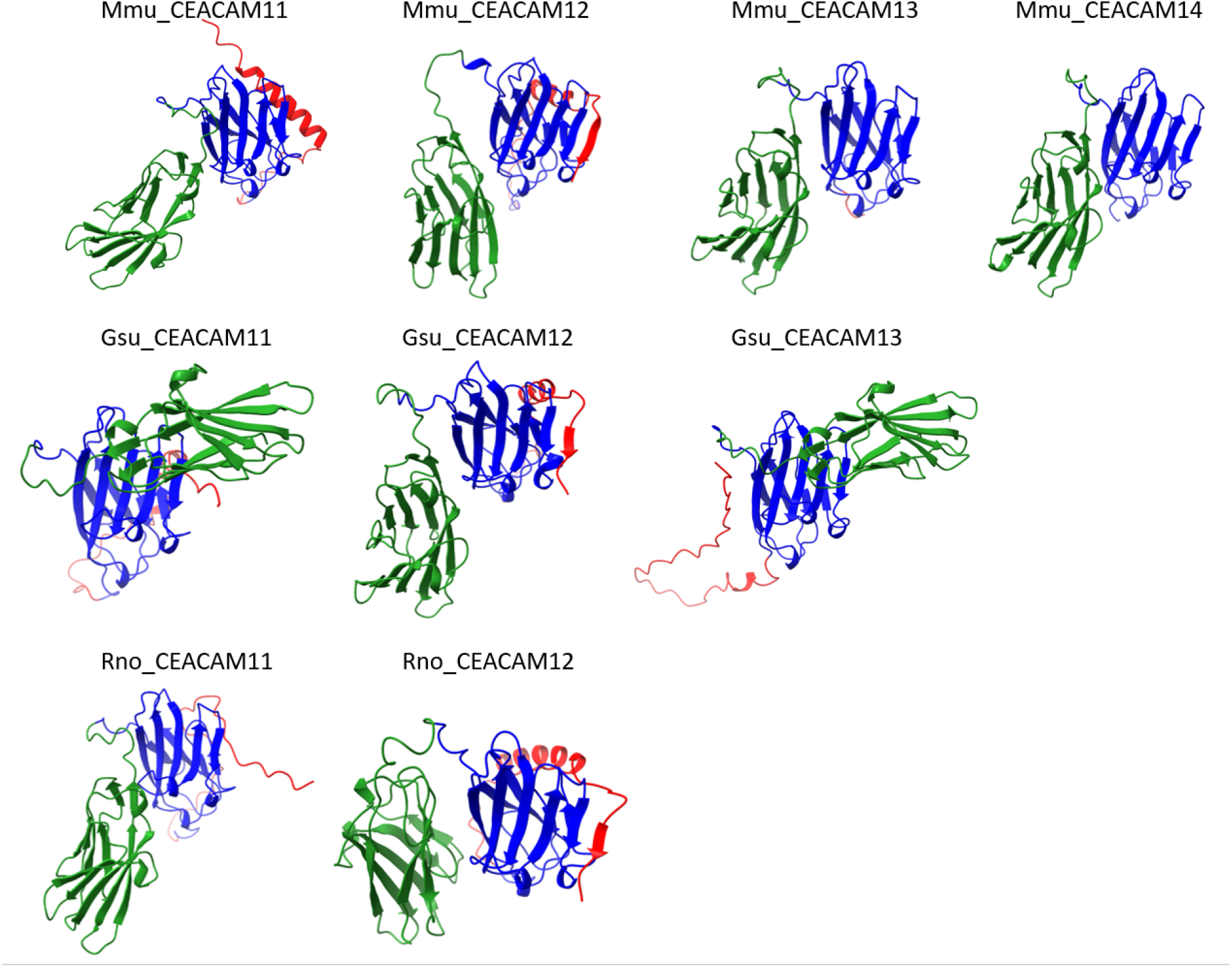
Structure of the expanded CEACAM11-related CEACAMs. The structure of the CEACAM11-related CEACAMs from mouse (Mmu), African thicket rat (Gsu) and rat (Rno) was predicted using ColabFold. The N1 domains are shown in green the N2 domains in blue and the exon 4-encoded domain in red. The arrows represent β-strands.

### *Ceacam11-14* genes are preferentially expressed in trophoblast cells

PSGs are defined as CEACAM1-related CEACAMs that are secreted and preferentially expressed in trophoblast cells [25]. Previously, we found that murine *Ceacam11-14* are expressed in placental tissues in the mouse [7]. Here we substantiated these findings by additional analyses of publicly available data sets as described in “Material and Methods”. Genes of the *Cea* gene family that were preferentially expressed in the placenta include *Ceacam9, Ceacam11-14*, and the *Psg* genes as determined by bulk mRNAseq data (Fig. 7A). scRNAseq data revealed that each of these genes is preferentially but not exclusively expressed by trophoblast cells in mice (Fig. 7B). In particular, *Ceacam14, Psg21, Psg23, Psg27*, and *Psg30* are expressed by additional tissue compartments in the placenta (Fig. 7B).

**Figure 7:**
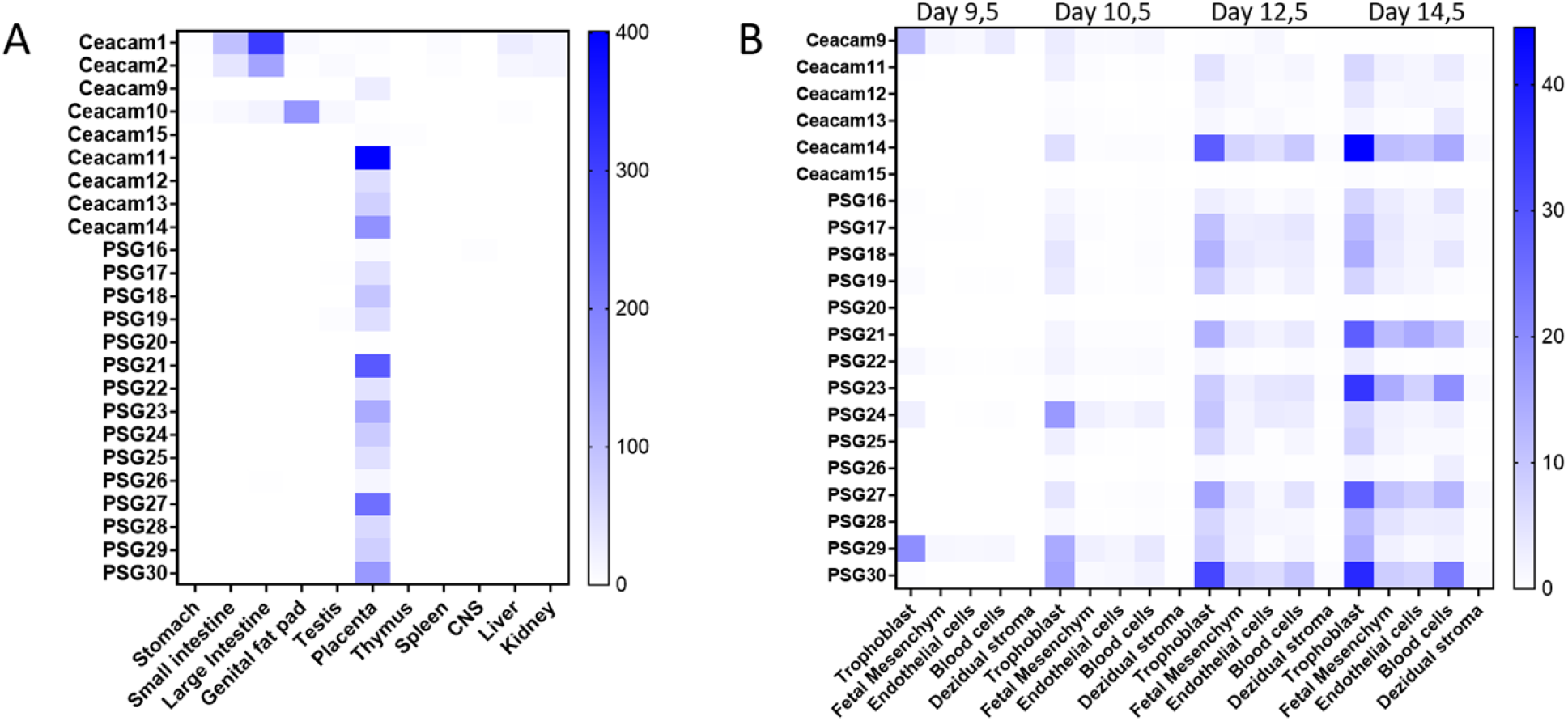
Expression of murine *Cea* gene family members. (A) Expression of murine *Cea* gene family members in different tissues as extracted from the Mouse ENCODE project (NCBI Geo BioProject: PRJNA66167). The relative expression is based on bulk mRNAseq and indicated by the color code as depicted on the right. (B) Expression of placenta-specific *Cea* gene family members by different placental cell types as determined by scRNAseq [26]. The relative expression is indicated by the color code as depicted on the right.

We further analyzed the expression of murine *Psgs* and *Psg-like Ceacams* by different trophoblast cell types at day 9.5, 10.5, 12.5 and 14.5 of pregnancy at single cell resolution. Overall murine *Psgs* and *Psg-like Ceacam* genes have a diverse expression pattern, although most genes were preferentially expressed by spongiotrophoblast cells and their precursors. However, in particular *Ceacam9* and *Psg29* were also expressed by glycogen cells. In addition, a significant expression of most *Psgs* and *Psg-like Ceacams* in syncytiotrophoblast cells and their precursors was noticed. *Psg23* showed the broadest expression pattern being expressed in different trophoblast cell types. *Ceacam15, Psg20, Psg22* and *Psg26* showed only a weak expression in placental cells at the investigated developmental stages. The expression of the majority of *Psgs* increased during pregnancy. In contrast, *Ceacam9* and *Psg29* showed the highest expression on day 9.5 followed by a decrease of expression. *Psg24* reached a peak of expression on day 10.5 (Fig. 8). *Ceacam11-14* showed a very similar expression pattern although with significant differences of expression intensities at the mRNA level (Fig. 8). Together this expression analyses strongly indicate that all *Ceacam*/*Psg* genes at the “rodent Psg locus” have to be consider as functional *Psgs*.

**Figure 8:**
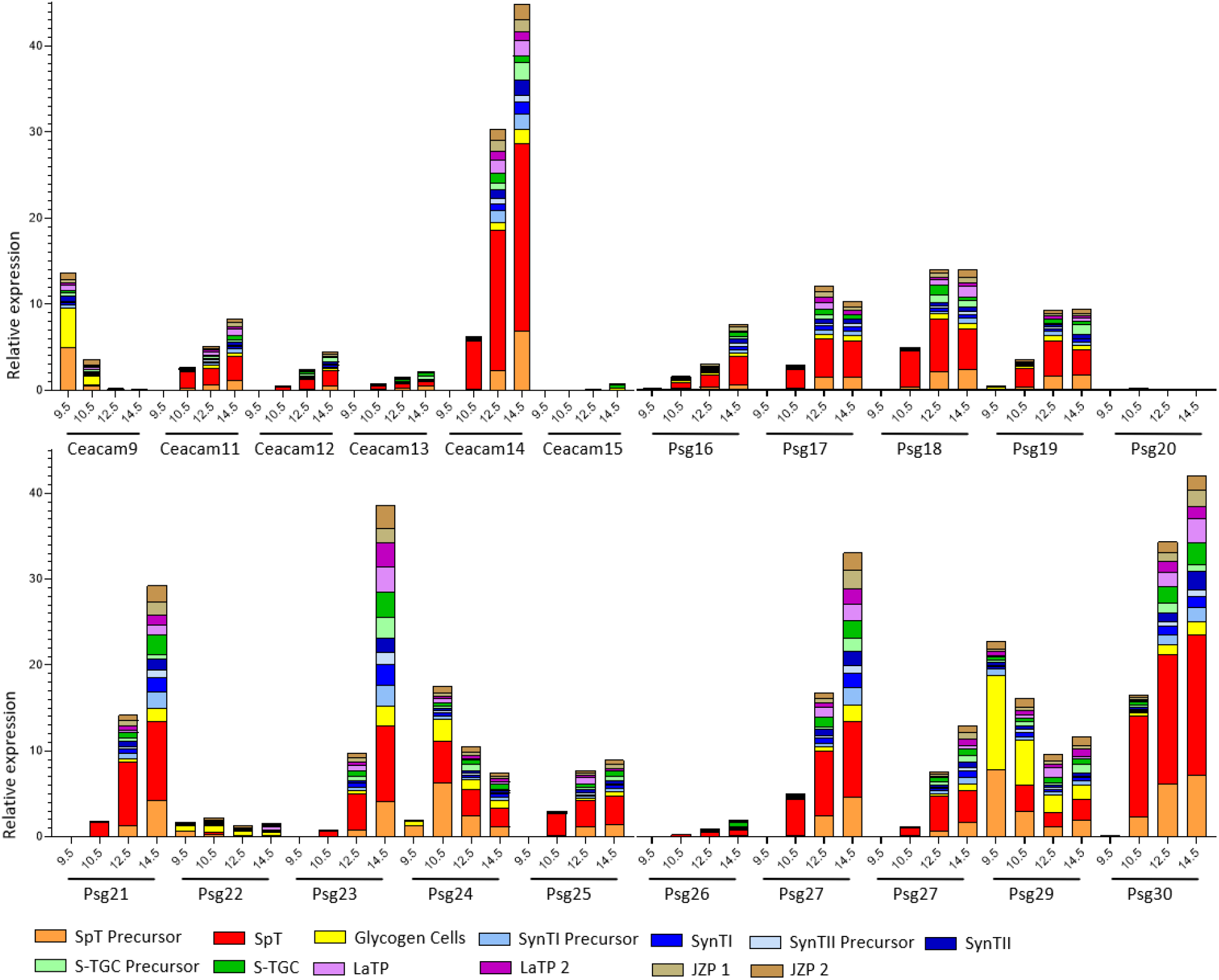
Relative expression of *Cea* gene family members in trophoblast subpopulations. The color code that identifies the different cell populations is shown below the graph. SpT, Spongiotrophoblast; SynT1, outermost syncytiotrophoblast layer, SynT2, syncytiotrophoblast layer between SynT1 and the fetal endothelium; S-TGC, sinusoidal trophoblast giant cells; LaTP, labyrinth trophoblast progenitor; JZP, Junctional zone precursor [26].

### The evolution of *Psgs* in Muroidea is highly dynamic

Variation of *Psg* copy numbers within Muroidea indicates a highly dynamic evolution of *Psg* genes in the *Psg* gene locus. However, there are significant differences between groups of *Psg*/*Psg*-like *Ceacam* genes. *Ceacam9* and *Ceacam15* are well-conserved single-copy genes. *Ceacam9* is found in all *Psg*-containing species. In some closely related Muroidea species (*M. ungulates, P. obesus, R. opimus*) *Ceacam9* appears to be a pseudogene due to a common 2 bp deletion in the N exons (Fig. 2; data not shown). In contrast, *Ceacam15* has been lost in the entire Arvicolinae subfamily (only *Ceacam15* gene remnants can be found in some Arvicolinae species: Elu, Eta, Mgl, Ozi). *Ceacam11* has been conserved for a certain time during which no amplification occurred. Only recently this gene has been amplified in Murinae. The *bona fide Psg* genes have been subject to multiple rounds of gene duplications and exon shuffling. Interestingly, gene expansion (possibly followed by gene loss in some groups of species) happened differentially at different subregions of the *Psg* locus of Muroidea species. While the number of *Psg*-like genes varies little in the Psg subregion flanked by the marker genes *Hif3a* and *Mill1* (9-12 *Psg* and *Ceacam11-14* genes), there is a large variation in *Psg* gene numbers in the *Psg* subregion flanked by *Mill1* and *Pglyrp1*, where between 1 (*M. coucha*) and 11 Psgs (*M. musculus*) are found (Fig. 9). In contrast, most of *Psg* gene size expansion by exon duplications occurred at the *Hif3a*/*Mill1* subregion (Fig. 9). Taken together, this complex evolutionary history makes the assignment of orthologous genes almost impossible between different families of Muroidea.

**Figure 9.**
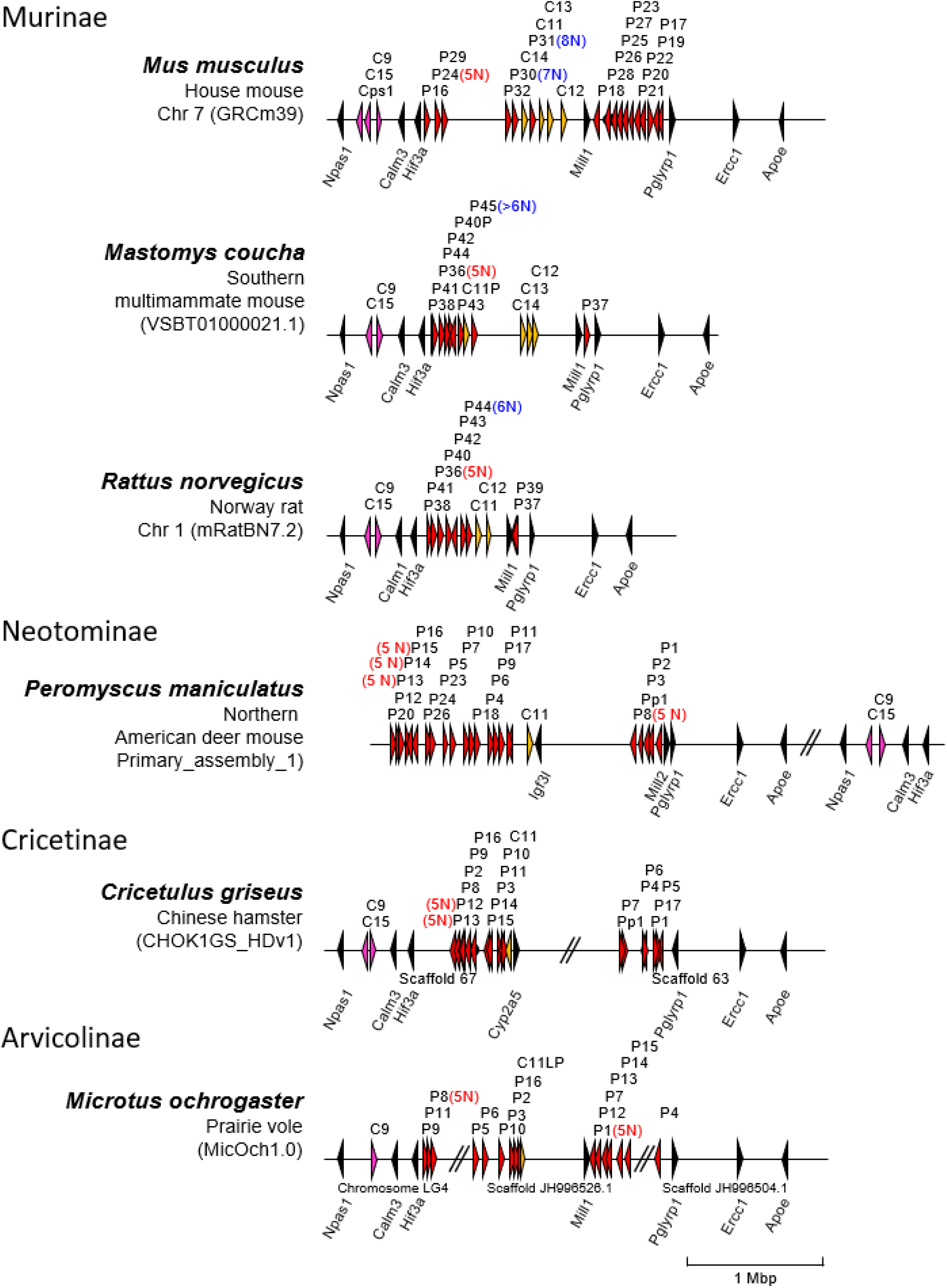
Divergent evolution of the *Psg* locus in Muroidea. The chromosomal arrangements of *Psg* genes at the *Npas1*/*Pglyrp1* locus of selected species of the Muroidea clade are shown. Subfamily and species names are shown at the left side. Arrowheads indicate genes with their transcriptional orientation. The *Psg*-related genes are shown in red (*Psg*), purple (*Ceacam9, Ceacam15* and *Ceacam pseudogene 1, Cps1*) or orange (*Ceacam11-14*), and selected flanking genes in black. The number of IgV domain-encoding exons exceeding the standard number 3 is indicated in brackets next to the gene names. IgV variant order as found in Mmu_Psg24 and Asp_Psg31 are shown in red and blue color, respectively. The *Psg* gene loci were aligned along the position of the *Npas1* gene. Databases and their versions used are listed below the species name. The borders of scaffolds are indicated by double slashes, their names below the chromosome. Of note: ortholog assignment of *Ceacam11-14* genes between species is not possible due to lack of unequivocal synteny and sequence relationship. Therefore, same gene names do not imply an orthologous relationship. Asp, Apodemus speciosus; C, Ceacam; Cps, Ceacam pseudogene; Mbp, million base pairs; Mmu, Mus musculus; P, pregnancy-specific glycoprotein (*Psg*) genes.

### The structure of PSG/PSG-like CEACAMs in rodents

Two principle domain compositions of PSGs/PSG-like CEACAMs were found in rodents, one group consist of two N domains and the other is built by one A domain and a variable number of N domains. Intact PSG-like CEACAMs built of two N domains are absent in various groups of rodents, including Nesomyidae, Avricolinae, and Gerbillinae (Fig. 10). The dominant domain composition of rodent PSGs is three N domains combined with one A domain (some 85 %), followed by PSGs comprising five N domains and one A domain (Fig. 10). Nevertheless, in each species analyzed at least one member is composed of one N domain and one A domain (Fig. 10).

**Figure 10:**
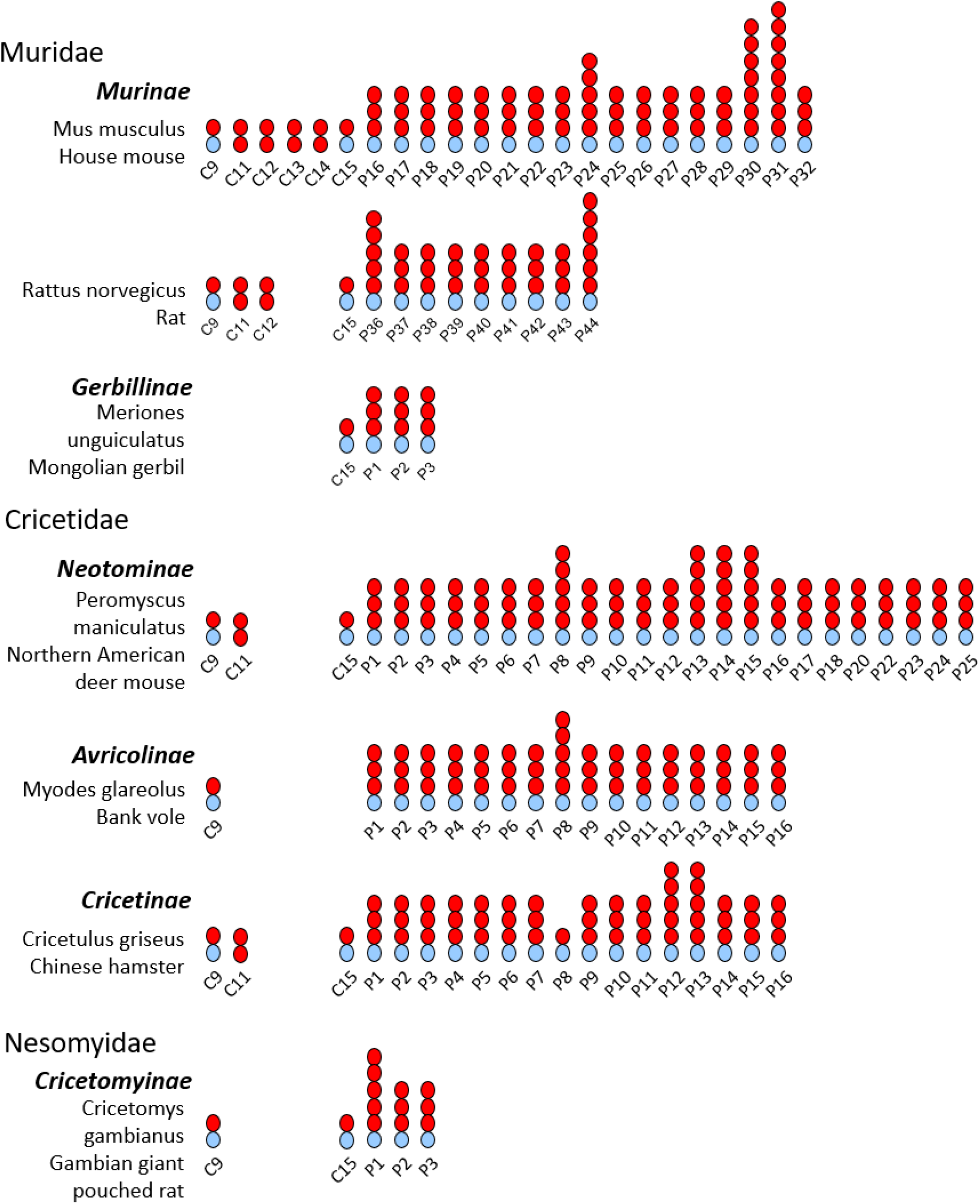
Domain structure of rodent PSGs/PSG-like CEACAMs. The domain organization of PSG/PSG-like CEACAMs from selected rodent species from the Muridae, Cricetidae, and Nesomyidae families was predicted by gene analysis. Family, subfamily and species names are indicated at the left side. Mouse and rat PSG domain organizations were confirmed by EST sequences when available. IgV-like domains are shown as red, and IgC-like domains as blue ovals. Note the highly variable number of PSG in the different rodent species (between 3 and 23). Identical PSG numbering does not imply an orthologous relationship. C, CEACAM; P, PSG.

### Variable evolutionary selection on individual genes and rodent populations

Previously we found that *PSGs* in bats after amplification are under selection for diversification. In primates, we observed a largely variable selection pattern depending on the species and domain examined. Here, we selected closely related groups of rodents where an orthologous relationship between genes could be identified and performed dS/dN analyses. Three rodent subfamilies could be analyzed Murinae, Neotominae, and Arvicolinae (Fig. 11). In all groups we found that *Ceacam9* is highly conserved i.e., under purifying selection (dN/dS < 1) mostly even more than the conserved *Ceacam19* gene (Fig 11 C, F, I). In contrast, the N domain of *Ceacam1* is under selection for diversification (positive selection) in Murinae and Arvicolinae while it is under purifying selection (negative selection) in Neotominae (Fig. 11 C, F, I) indicated by dN/dS values >1 and <1, respectively. Remarkably, the positive selection of the *Ceacam1* N domain exon in Murinae as in other species (e.g. humans) is thought to be the result of pathogen usage of CEACAM1 as an entry receptor. In general, PSGs in rodents are under negative selection (Fig. 11 A, B, D, E, G, H). In Neotominae and Arvicolinae individual N domains and A domains show a relaxation of negative selection. In particular, the N2 domains of PSG8 and PSG11 in Neotominae exhibit or are close to positive selection, respectively. In Arvicolinae several N and A domain exons show a relaxed negative selection (0.5 < dN/dS < 1.0) (Fig. 11 G, H). The single *CEACAM11-14* gene in Neotominae is under negative selection (dN/dS = ~ 0.4), in contrast in Murinae the *Ceacam11-14* genes show some relaxation of purifying selection, indicating that upon amplification the newly generated genes underwent some adaptation to their new functions Fig. 11 C.

**Figure 11.**
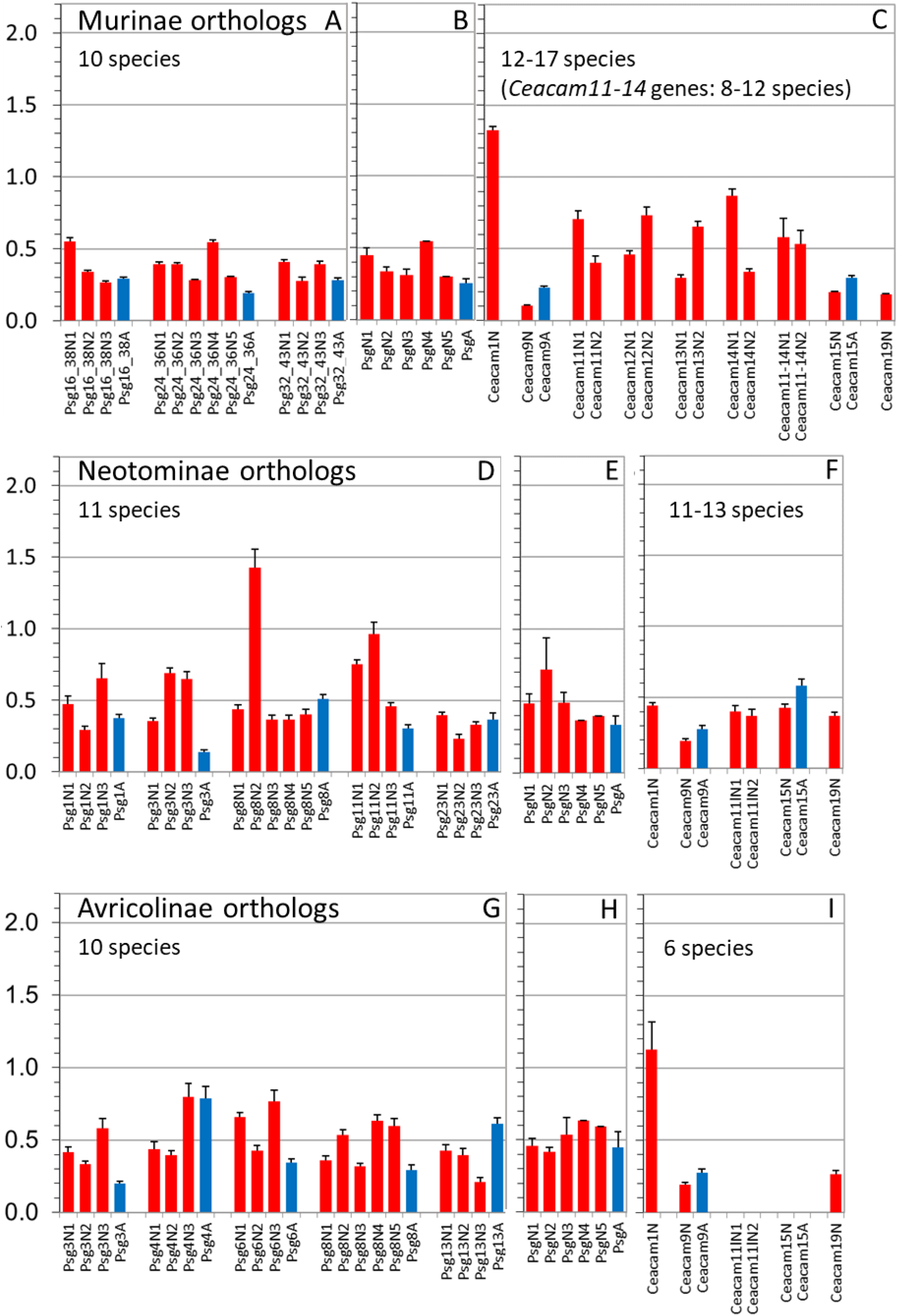
Differential selection for diversification in Ig domain exons of trophoblast-specific *Ceacam*/*Psg* genes. Nucleotide sequences of N and A domain exons of *Psg* orthologs from rodent species A-C: Murinae (Ani, Asy, Gsu, Hal, Mca, Mco, Mmi, Mmu, Mmu_cas, Mmu_mus, Mna, Mpa, Msi, Msp, Pde, Rdi, Rno, Rra), D: Neotominae (Nle, Oto, Pat, Paz, Pca, Per, Ple, Pma, Pme, Pnu, Ppo) and G: Arvicolinae subfamilies (Elu, Eta, Mag, Mar, Mfo, Mgl, Moc, Moe, Mor, Ozi) were compared pair-wise in all combinations after manual removal of gaps and the ratio of the rate of nonsynonymous (dN) and synonymous mutations (dS) was calculated for whole N and A domain exons and the mean ratios were plotted. The whiskers represent standard errors of the mean (SEM). Only genes were included for which an orthologous relationship could be demonstrated by phylogenetic analyses using CLUSTALW. The dN/dS values of three to five PSG genes for each exon type and subfamily were averaged and plotted (B, E, H). In addition, the dN/dS ratios for the N domain exons of trophoblast-specific *Ceacam* genes and, for comparison, for *Ceacam1* and *Ceacam19* orthologs of the same rodent species were calculated (C, F, I). These two genes are known to represent genes under diversifying (dN/dS > 1) and purifying selection (dN/dS <<1), respectively, in other mammalian species [23]. Of note: In Neotominae species, for the single gene related to the CEACAM11-14 genes in Murinae no ortholog could be clearly identified. A, IgC-like domain exon; N, N domain exon; CEACAM11L, CEACAM11-like. For the three letter species name abbreviations please refer to Supplementary Table 1.

### Independent evolution of *Psgs* in rodents without a “rodent *Psg* locus”?

Since structurally different *Psgs* evolved in Muroidea it is worth speculating that in other rodents *Psgs* evolved at a different locus in the genome as found for Muroidea. Indeed, we found some amplification of *Ceacams* at the *Ceacam* locus flanked by the marker genes *Cd79a* and *Xrcc1*. However, there is no evidence that these genes represent *bona fide* genes that encode secreted proteins or are expressed in a trophoblast-specific manner. In contrast, in species where we further analyzed the expanded *Ceacams* we found also an expansion of transmembrane domain coding exons, suggesting that these are membrane-bound *Ceacams*. Nevertheless, since expression data are lacking we cannot exclude that *Psgs* evolved in other rodents than the ones described in the present report.

## Discussion

*PSGs* were so far described in primates, mice and rats, microbats, and the horse [9, 14, 16]. With the exception of the horse, these species have a hemochorial placenta. Thus, we have previously speculated that the intimate contact of trophoblast cells with maternal immune cells drives the evolution of *PSGs* [11, 16, 23]. Indeed, in primates, the emergence of *PSGs* correlates with the appearance of hemochorial placentation [16]. The only primates so far identified that have a hemochorial placenta but no *PSG*s are the tarsiers [16] indicating that in primates *PSG*s evolve almost in parallel to a hemochorial type of placentation. However, while the amplification of *PSG* genes in New World monkeys remained limited (1-7 *PSG* genes) a massive amplification occurred in Old World monkeys resulting in more than 20 gene copies in some species [16]. These differences may be due to unknown restrictions of successful gene duplication at the *PSG* locus in New World monkeys or by a relaxed selection pressure for *PSG* gene amplification. In order to get further insights into the evolution of *PSG* genes we analyzed the evolution of *Psg*s in rodents. Since all rodents, with very few exceptions, have a hemochorial placenta we expected that in most if not all rodents *Psgs* are present, although they have been only described in mice and rats so far. It has been suggested that the common ancestor of rodents had a hemochorial placentation with an interhemal barrier that had a single layer of syncytial trophoblast cells [18]. This anatomical feature was retained in the clade comprising Hystricomorpha (guinea pigs and others) and Sciuromorpha (squirrels) [18]. In contrast, in Myomorpha (mice and others) several placental transformations occurred [18], most remarkably within the Muridae family, which has a special three-layered trophoblast [27, 28]. The three-layered trophoblast containing a layer of cytotrophoblast and two layers of syncytiotrophoblast cells appeared together with the capture of the *syncytin-A* and *syncytin-B* genes in the most recent common ancestor (MRCA) of Muroidea including Muridae, Cricetidae, and Spalacidae family species [29]. Of them, the Spalacidae is the only family in which *Psgs* did not evolve indicating that shortly after the invention of the three-layered trophoblast *Psgs* evolved. This may refine our picture of the forces driving PSG development. It may be that alterations of the fetomaternal interface create opportunities to optimize the molecular fetomaternal crosstalk. Members of the CEA family may be predisposed to fulfill this task once they are secreted by fetal trophoblast cells. Such a “beneficial” *PSG* gene may then be fixed in the genome and eventually amplified. Because the fetomaternal interface evolves extraordinarily fast such changes may frequently occur thus explaining why PSGs can evolve independently multiple times in different mammalian lineages. Since *Ceacam9, Ceacam15*, and *Ceacam11-like* genes or at least remnants of the latter are present in the genome of *Psg*-harboring rodents it is not possible to decide which ancestor of these genes is the primordial gene of rodent *Psgs*. However, a combination of *Ceacam9* or *Ceacam15* with *Ceacam11-14* would provide all building blocks (three N domain exons and one A domain exon) to create typical rodent *Psgs*. The strong correlation between the existence of *Psgs* and the presents of *Ceacam9* may indicate that *Ceacam9* plays a pivotal role in the evolution of *Psg* genes. If *Ceacam9* is the founder of *Psgs, Ceacam15* may be an early duplicate of *Ceacam9* which gained a new function but was not further amplified. The high conservation of *Ceacam15* argues for such a speculation. On the other hand, *Ceacam15* and the ancestor of Ceacam11-14 were lost in Arvicolinae indicating that in the MRCA of this group, both genes lost their function and therefore were subsequently deleted from the genome. Rodent PSGs are in general composed of three (more rarely of five, six, seven or eight) IgV-like domains and one A domain of the A2 type. Since the vast majority of rodent PSGs are composed of the typical exon arrangement with 3 exons coding for IgV-like domains and one IgC-like domain we conclude that once a Psg gene had evolved the duplication of whole Psg genes was the major mechanism of Psg gene amplification in rodents. The expansion of PSGs is still ongoing as indicated by the different number of *Psg* genes and their independent expansion e.g. in mice and rats. In addition, as previously shown for mouse *Psgs, Psgs* of other Muroidea evolve extremely fast therefore orthologs can only be assigned between very closely related species (Fig. 4) [24]. The fast evolution limits the possibility to analyze the nature of selection on rodent *Psgs* (Fig. 11). Nevertheless, our results indicate that some PSGs in some species are under positive selection, but the majority are under purifying selection. These results suggest that most rodent PSGs have adapted to a certain function while only some, possibly newly duplicated, PSGs are free to acquire novel functions or ligands. More recently, a second wave of gene amplification took place. The ancestor of *Ceacam11-14* is under purifying selection in all species that have only one gene. In Murinae the purifying selection seems to be relaxed, enabling some flexibility for functional optimization (Fig. 11). Remarkably, CEACAM11-14 are structurally different from the bona fide PSGs in rodents composed of only one N domain and one A domain. However, the very similar expression pattern of *Ceacam11-14* and *Psgs* in placental cells (Fig. 7; Fig. 8; [30]) suggest that both are functional “PSGs”. We have previously reported that PSGs are structurally different in different species, due to an independent evolution. This is now the first report showing that PSGs did evolve twice in one mammalian group, leading to structurally distinct PSGs. This indicates that the birth of PSGs is a frequent event explaining the independent evolution in various mammalian lineages.

Since the translocation of a *Ceacam* gene family member or parts of it seem to be a hallmark of the evolution of *Psgs* in rodents the question arises what kind of *Ceacam* gene was translocated to form the original *Psg* locus? One possibility can be envisaged that part of an ITAM-containing *Ceacam* gene was translocated with concomitant destruction/loss of the ITAM motif-encoding region of the gene. Such a scenario would explain the strong correlation between the absence of ITAM-containing CEACAMs and the presence of PSGs (Fig. 3). In rodents without PSGs, ITAM-containing CEACAMs exist, as in most other mammalian species (Fig. 3) [23]. Thus, this report shows for the first time that most rodents have ITAM-harboring CEACAMs and that the loss of ITAM-containing CEACAMs happened only recently affecting the species of the Muridae family. A summary of the possible evolution of *Psgs* in rodents is depicted in Supplementary Figure 2.

Although we did not find any evidence for the presence of PSG in other rodents we cannot exclude that they may exist in some species due to their structural variability and missing expression data of most species analyzed in this report. In addition, we are aware that the simplified construction of rodent phylogeny used in this study by comparing the IgV-like (N) domain exons of *CEACAM19* did not completely mirror the previously published studies using more complex molecular data [21, 22, 31]. In contrast to these studies, we did not see a monophyletic clade comprising Hystricomorpha and Sciuromorpha. In addition, the Castorimorpha did not appear to be a sister group of the Myomorpha as previously shown. Nevertheless, the relationship between the Muridae and Dipodidae as well as the relationship within the Muridae family agrees with published data [21, 22, 31].

In summary, the expansion of the analysis of the CEA gene family to the entire rodent clade shed new light on the evolution of the CEA gene family of the most frequently used animal models for medical research, i.e. mice and rats. This study demonstrates that the loss of an ITAM-encoding *Ceacam* gene and the appearance of *Psg* genes is a rather recent event in rodents only affecting the Cricetidae, Muridae and Nesomyidae families.

## Methods

### Identification and nomenclature of genes

Nucleotide and amino acid sequence searches were performed using the NCBI BLASTBLAT tools (http://www.ncbi.nlm.nih.gov/BLAST) and the Ensembl database (http://www.ensembl.org/Multi/Tools/Blast?db=core) using default parameters. For the identification of rodent *Ceacam* exons, *Ceacam* and *Psg* exon and cDNA sequences from known mouse and rat *Ceacam*/*Psgs* were used to search various databases at NCBI and Ensemble including whole-genome shotgun contigs (wgs), and Transcriptome Shotgun Assembly (TSA). A comprehensive overview of the used genomic data sources for the analyzed rodent species is given in Supplementary Table 1. Hits were considered to be significant if the E-value was < e^-10^ and the query cover was > 50%. Genes that contained stop codons within their N domain exons or lacked appropriate splice acceptor and donor sites in these exons were considered to represent pseudogenes. Nucleotide sequences from the N domain exons can be used as gene identifiers (Supplementary File 2). The same strategy was employed to identify other genes of the CEACAM families. *Ceacam* genes, the N exons of which exhibited >99% nucleotide sequence identity, were considered to represent alleles.

### Quantification of PSG expression

For the quantification of murine PSG expression, we reanalyzed publicly available datasets, these include mRNA sequencing data sets generated by the Mouse ENCODE project available at NCBI Geo BioProject: PRJNA66167 as well as single cell mRNA sequencing data available at https://figshare.com/projects/Single_nuclei_RNA-seq_of_mouse_placental_labyrinth_development/92354 [26, 32].

### Sequence motif identification and 3D modeling

The presence of immunoreceptor tyrosine-based activation motifs (ITAM), ITAM-like, and immunoreceptor tyrosine-based inhibition motifs (ITIM) and immunoreceptor tyrosine-based switch motifs (ITSM) were confirmed using the amino acid sequence pattern search program ELM (http://elm.eu.org/). Transmembrane regions, and leader peptide sequences were identified using the TMHMM (http://www.cbs.dtu.dk/services/TMHMM-2.0/), the SignalP 4.1 programs (http://www.cbs.dtu.dk/services/SignalP/), respectively. The structure predictions of murine CEACAM11-14 and rat CEACAM11-12 were retrieved from the “AlphaFold Protein Structure Database”. The structure of CEACAM11-13 from African tree rat (Grammomys surdaster) was predicted using “ColabFold” [33].

### Phylogenetic analyses and determination of positive and purifying selection

Phylogenetic analyses based on nucleotide and amino acid sequences were conducted using MEGAX [34]. Sequence alignments were performed using Muscle implemented in MEGAX. Phylogenetic trees were constructed using the maximum likelihood (ML) method with bootstrap testing (500 replicates) and the Tamura-Nei substitution model [35]. Other multiple sequence alignments were performed with CLUSTALW programs (http://npsa-pbil.ibcp.fr/cgi-bin/npsa_automat.pl?page=/NPSA/npsa_clustalw.html;http://www.genome.jp/tools/clustalw/). In order to determine the selective pressure on the maintenance of the nucleotide sequences, the number of nonsynonymous nucleotide substitution per nonsynonymous site (dN) and the number of synonymous substitutions per synonymous site (dS) were determined for *Psg* and *Ceacam* N domain and IgC-like exons. The dN/dS ratios between pairs of *Psg* orthologs and paralogs and orthologous *Ceacam* genes were calculated after manual editing of sequence gaps or insertions guided by the amino acid sequences using the SNAP program (Synonymous Nonsynonymous Analysis Program; http://www.hiv.lanl.gov/content/sequence/SNAP/SNAP.html) [36].

## Declarations

### Ethics approval and consent to participate

Not applicable

### Consent for publication

Not applicable

### Availability of data and materials

All relevant data are publicly available and described in the manuscript

### Authors’ Contributions

R.K. conceived the study, carried out data analysis and drafted the manuscript. W.Z. performed most of the data mining and contributed equally to manuscript writing. Both authors read and approved the final manuscript.

## Acknowledgements

Not applicable

